# Rich Experience Boosts Functional Connectome and High-Dimensional Coding in Hippocampal Network

**DOI:** 10.1101/2022.02.23.480123

**Authors:** Brett Addison Emery, Xin Hu, Shahrukh Khanzada, Gerd Kempermann, Hayder Amin

## Abstract

Challenging the brain with experiential richness creates tissue-level changes and synaptic plasticity, but the interjacent network level has not been accessible. We here show that environmental enrichment has unexpectedly far-reaching effects on network connectivity and multi-dimensional coding in the hippocampus. We present direct evidence that experience impacts local and global network connectivity, synchrony, and rhythmic dynamics. For this, we investigated the hippocampi from standard-housed mice (SD) and mice living in an enriched environment (ENR) using large-scale *ex vivo* recordings with a high-density microelectrode sensing array that – with the unprecedented spatiotemporal resolution–allowed simultaneous electrophysiological assessment across the entire circuit. In the absence of extrinsic electrical network stimulation, we found enhanced functional connectivity and high-dimensional coding in hippocampal-cortical networks of ENR mice. The mapped connectome illustrated a scale-free smallworld topology and an ENR-induced resilience to random failures. ENR enhanced large-scale spatiotemporal firing patterns, which facilitated efficient pattern separation and boosted the information encoded in the firing phases of slow oscillatory rhythms. Given that essentially all electrophysiological studies on network behaviors have been done on animals housed in stimulus-poor conditions, our SD mice showed the expected normal functionality. The literature consequently underestimates the extent of spontaneous network activity and connectivity under truly physiological conditions. Our results pave the way to unveil fundamental mechanisms of experience-dependent enhancement in the hippocampal network underlying high brain functions and provide markers for large-scale network remodeling and metaplasticity.

## INTRODUCTION

While the classical experimental paradigm of “environmental enrichment” has led to a large body of evidence about how environmental stimuli affect the brain^1–3^, knowledge about large-scale effects on the network and circuit level has been scarce^4^. Nevertheless, such effects have been generally implicated in the notion that the fine structure of the brain is dependent on past activity^4^. This lack of direct evidence at the circuit level is partly explained by the historical lack of applicable methodology. Broad-scale electrophysiology has been impossible and non-invasive MRI in humans suggests that enrichment has generalizing network effects^5^ does not have sufficient resolution to study particular brain networks in rodents^6^. The hippocampal network plays a central role in spatial, contextual, and episodic memory, as well as effective behaviors, and the hippocampus is sensitive to environmental influences that result in functional changes in these domains^7,8^. However, the connectivity framework and the computation for this network plasticity have largely remained elusive. Electrophysiological studies comparing ENR with SD brain in rodents indicated enrichment effects such as increased LTP in single concrete neuronal systems^9^ but with their low number of cells that are addressable at the same time could not generalize to the network level of the hippocampus. They could not answer the fundamental question of whether ENR would promote refinement of the connectome and result in increased and more complex oscillatory firing patterns. Functional MRI studies in humans have conversely indicated that decreased resting-state activity, present in the absence of external stimuli, is associated with impaired brain function in aging and neurodegeneration^10^. To thus address the fundamental question of how exposure to richer environmental stimuli might affect neuronal network architecture, topology, and activity, we combined novel large-scale microelectrode arrays that allow us to simultaneously measure neuronal activity across the entire rodent hippocampal circuit with advanced computational analyses. The size of the array allowed us to characterize the hippocampal network in its cortical context.

## RESULTS AND DISCUSSION

We aimed at quantitatively mapping the spatiotemporal dynamic changes of large-scale functional hippocampal-parahippocampal cortical (Hippo-cortical) circuitry in mice reared in enriched environmental conditions (ENR) compared to standard-housing (SD). We examined fundamental questions of how exposure to ENR remodels the functional neuronal network architecture, topology, and coding, as these would explain the well-described enhancement of cognitive capabilities and learning in ENR. We therefore obtained next-generation electrophysiological data by recording simultaneous extracellular discharges with high-density microelectrode arrays **(Figures 1a** and **b**). We outlined the hippo-cortical circuit as six interconnected regions (DG, Hilus, CA3, CA1, entorhinal cortex ‘EC’, and perirhinal cortex ‘PC’) by overlaying the bioelectrical functional readouts with structural images.

**Figure 1.**
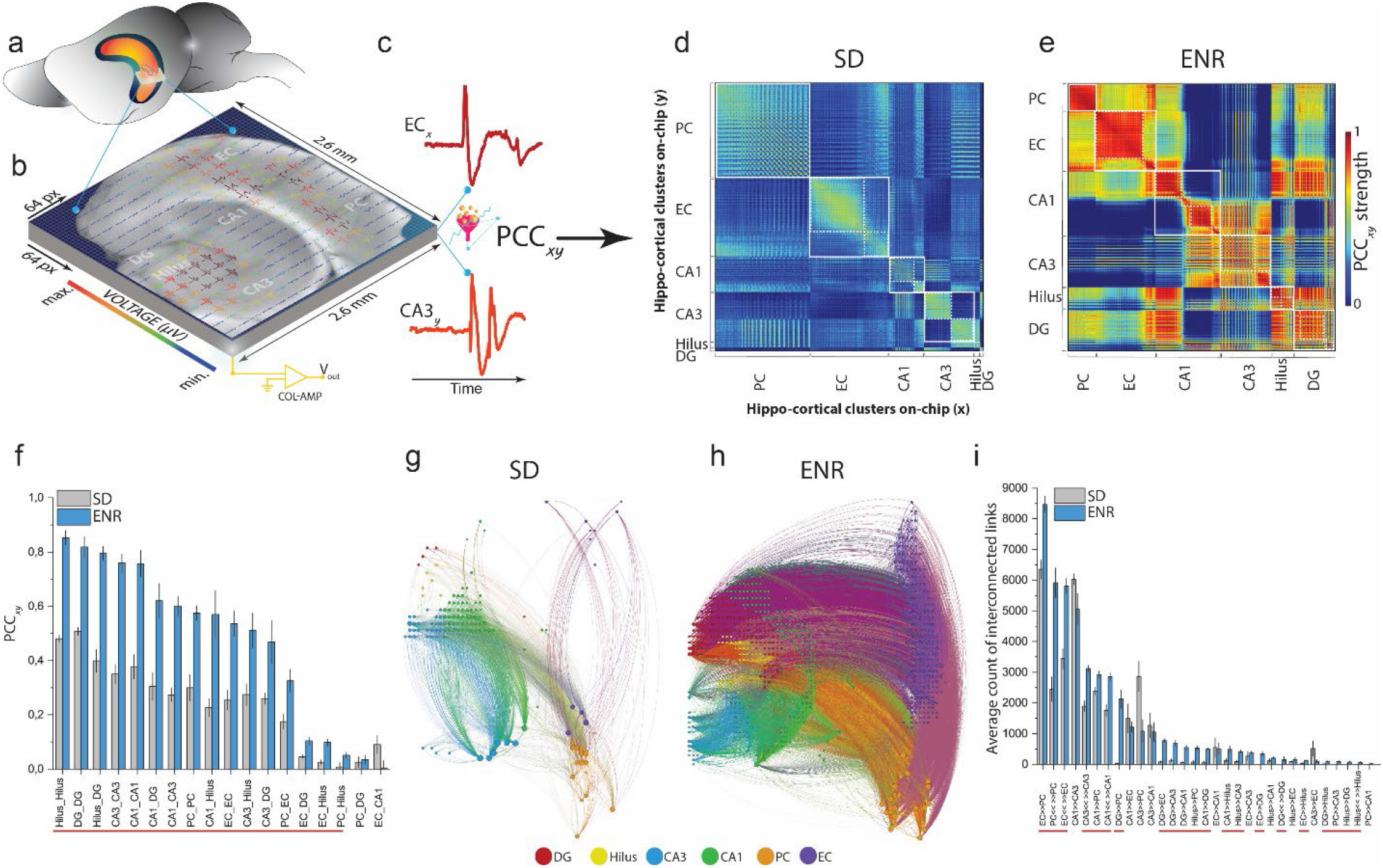
ENR promotes large-scale functional connectome in hippo-cortical network. **a)** Mouse brain drawing displays the hippocampus and a transverse section of the slice. **b)** The entire hippo-cortical slice is overlaid on the 64×64 electrode array allowing to monitor the 6 interconnected subregions simultaneously with their concurrent structural information. **c)** Exemplary pairwise firing electrodes in the chip array to extract time courses to compute the PCC. **d-e)** Functional connectivity matrices representing cross-correlation of SD and ENR networks constructed and clustered by the pairs of firing neurons in their associated interconnected subnetworks. ENR displays higher correlation values in all hippo-cortical networks. White boxes located along the diagonal illustrate modular graphs, where a set of nodes are highly connected in a region, while white dotted boxes illustrate hierarchical modular graphs that are submodules of the large modules. **f)** Quantification of the correlation matrices shows a significant increase of PCC in interconnected regions of the ENR network compared to SD. Red line indicates the interconnected subnetworks with significance differences (\**p < 0.05*, \*\**p < 0.01 ANOVA*) (n=48 slices from 6 mice of SD and ENR each). **g-h)** Connectome mapping of the hippo-cortical network in ENR and SD. The network visualization was done using the Gephi program 9.2 version (https://gephi.org) to illustrate 2% of the total connection in SD (node= 136, and links= 1762) and ENR (node= 661, and links= 21648) networks, which indicates a salient higher density of connection in ENR (h) compared to SD network (g). Graph nodes are scaled according to degree strength and colored according to hippo-cortical module association and indicated in colored circles legends. Colored links identify the intra- and intercluster connections. **i)** Quantification of the directed interconnected links shows higher unidirectional and bidirectional communication in the ENR compared to the SD network. Red line for significant interconnections (\**p < 0.05 ANOVA*) (n=48 slices from 6 mice of SD and ENR each).

### Enhanced functional connectivity and organization in ENR networks

In order to quantitatively characterize the bidirectional inter-regional relationship, we determined the firing patterns of concomitantly active neuronal ensembles by computing the cross-covariance of pairwise firing electrodes using Pearson’s correlation coefficient (PCC)^11^. Analysis of the slow oscillatory component of local field potentials (LFPs) up to 100 Hz yielded association matrices of firing coefficients with dimensions equal to the number of firing electrodes in the 64 x 64 chip array (see method). We found significant enhancement in the local and global strength of spatial interactions in the ENR network compared to SD, as shown in the cross-correlogram and the quantified differences between connectivity matrices of pooled individual slices from several preparations **(Figures 1d-f)**. Both SD and ENR networks displayed modular functional organization of hippo-cortical connectivity architecture that featured groups of nodes densely interconnected in clusters (i.e., white rectangles in **Figures 1d** and **e**). However, compared to SD, the ENR network featured a more hierarchical modular arrangement indicated by large modules consisting of smaller interconnected modules (i.e., white dotted rectangles in **Figures 1d** and **e**). This confirms strengthened local connectivity in the ENR group to optimize network rewiring, adaptability of function, and pattern evolvability without compromising the functional integrity of other network clusters^12^.

To estimate the mesoscale functional connectivity and measure the information flow and its direction within the correlated links in the circuitry, we employed multivariate Granger causality and directed transfer function (DTF)^11^. The ENR network showed both higher unidirectional (i.e., DG→CA3, CA1, EC, PC) and bidirectional (i.e., CA3↔CA3, CA1↔CA1, DG↔DG, Hilus↔Hilus, EC↔EC, and PC↔PC) interaction links compared to the SD network **(Figures 1g-i)**.

Next, to assess the impact of ENR on the spatial and topological complexity of large-scale connection patterns of the network, we employed graph-theoretic analysis to generate large-scale graphical metrics of local and global functional connectivity (**Figure 2a**). The network can be mathematically described as neuronal nodes linked by functional connections as edges. We computed the graph metrics from connectivity patterns of co-firing neuronal ensembles (corresponding to the firing electrodes of the array) encoded in the LFPs under SD and ENR conditions (**Figures 2b** and **c**).

**Figure 2.**
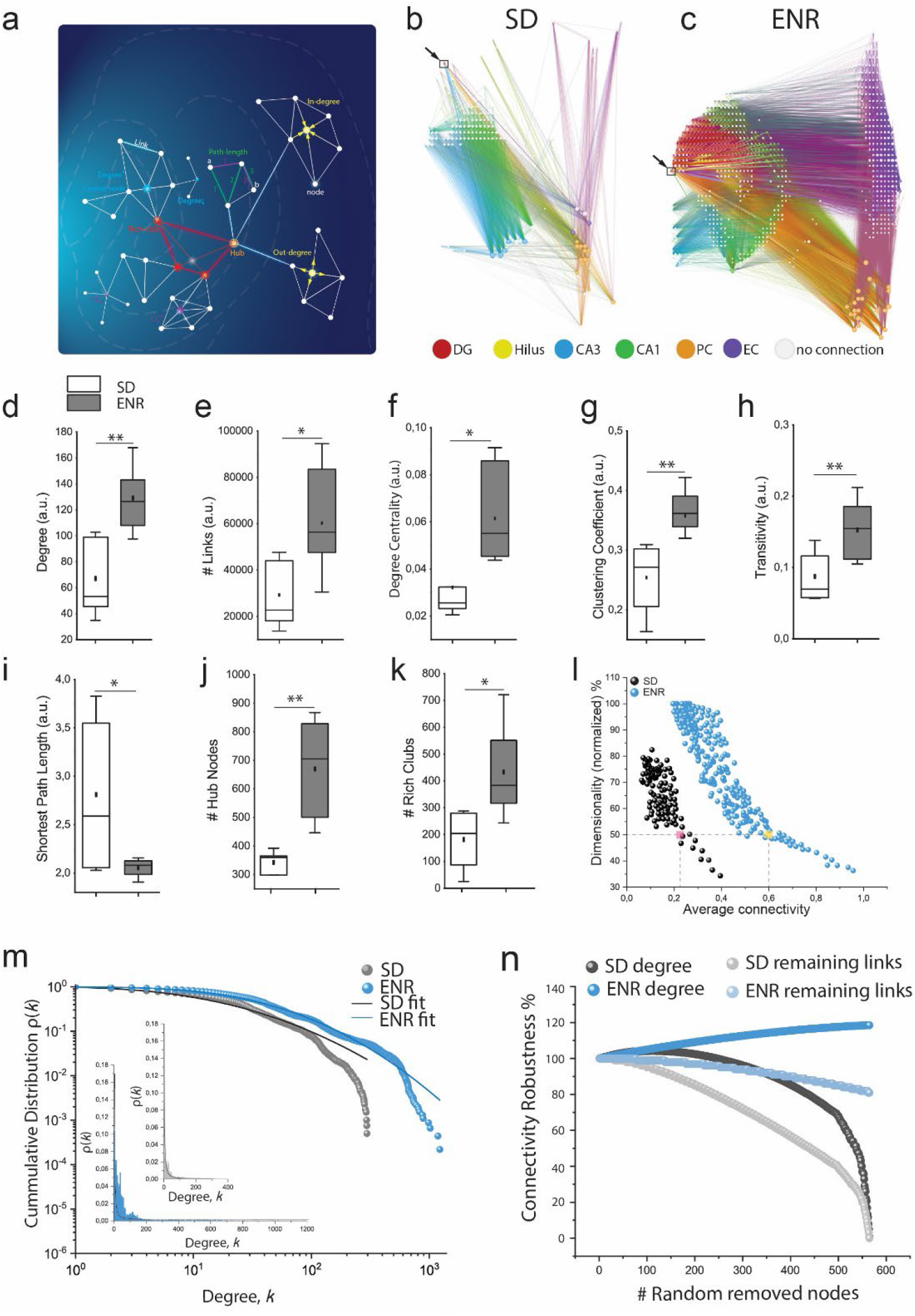
ENR boosts network complexity and coding dimensionality. **a)** Schematic illustration of key complex network metrics of large-scale hippo-cortical network on HD-chip array. **b-c)** Functional connectivity maps of the spatial distribution of interconnected subnetworks showing increased complexity of ENR compared to SD networks. Arrow indicates the exemplary selected node in the DG layer to identify the associated local and global interactions. **d)** ENR network exhibits higher complexity shown by higher node degree (\*\**p < 0.01 ANOVA*). **e-f)** ENR network is significantly densely interconnected compared to SD indicated by higher numbers of links and node degree centrality (\**p < 0.05 ANOVA*). **g-h)** Graph clustering coefficient and transitivity confer significant network functional segregation in ENR than SD (\*\**p < 0.01 ANOVA*). **i)** ENR networks showed lower path length indicating higher functional integration than SD networks (\**p < 0.05 ANOVA*). **j-k)** ENR networks display highly dense connected hub node and emerged rich club organization compared to SD networks (\*\**p < 0.01, *p < 0.05 ANOVA*). **l)** ENR network exhibit greater dimensional space for information coding than SD computed as a function of average connectivity (*p < 10^-8^ Kolmogorov-Smirnov test*). Although dimensionality is reduced to 50% (colored squares), ENR maintains higher connectivity than SD. **m)** Scale-free functional network topology with small-world properties in SD and ENR networks indicated by the power-law distribution. The log-log plot of the cumulative connection distribution for ENR networks exhibits a significantly heavier-tail than SD networks indicating low degree nodes to coexist with a few densely connected hub, yet higher than SD networks (inset), which reach a cutoff on the power-law lead to steep decay of the tail of connectivity distribution (\**p < 0.05 Kolmogorov-Smirnov test*). The power-law distribution was fitted by the lognormal function with goodness-of-fit in log-log plot (with a coefficient of determination R^2^ = 0.96 and 0.99 for SD and ENR, respectively). **n)** ENR networks display higher connectivity robustness to random failures quantified by the percentage of degree and remaining links after random node removal (SD vs. ENR degree and remaining links, *p < 10^-8^ Kolmogorov-Smirnov test*).

As revealed by the node degree analysis and its ingoing and outgoing connections, we detected a much greater network complexity in ENR compared to SD (**Figure 2d** and **Supplementary Figure 1**). We also found a significant increase in the functional interaction with other nodes in the ENR network conferred by the total number of links and degree centrality (**Figures 2e** and **f**). Computing the clustering-coefficient (CC) and transitivity of SD and ENR networks, we found that ENR networks were consistently more segregated than SD networks^13^ (**Figures 2g** and **h**). We also found significantly shorter paths in ENR compared to SD networks, indicating greater network integration in ENR^13^ (**Figure 2i**).

ENR networks exhibited more densely-interconnected nodes (i.e., hub nodes) and formed more ‘rich-club’ organizational structures than SD, indicating greater higher-order functional specialization, the resilience of the network, and increased capacity of global communication^14^ in ENR (**Figures 2j** and **k**).

Dimensionality of a network refers to the size of its neuronal population as well as to the dynamic range of the firing activity of its individual neurons. Hence, a high dimensional coding suggests powerful and flexible neural computations and endows neural circuits with a larger coding space to facilitate the separation of overlapping activity representations^15,16^. We found that as a function of the average connectivity, the ENR network exhibited higher dimensionality than the SD network and that in ENR with increasing connectivity (i.e., pairwise correlation of activity patterns), dimensionality of the coding space decreased more protractedly than in SD; thus endows the network with a greater capacity for linear separability of overlapping patterns^17^; (**Figure 2l** and **Supplementary Figure 2**).

Small-word topology is a remarkable feature of many biological and business networks^18^ that is characterized by a scale-free architecture with highly connected hub nodes and a degree distribution that decays with a power-law tail^19^. Scale-free networks provide error tolerance and resilience to random failures^20^. We found that the functional node degree distributions of SD and ENR networks followed a power-law function (i.e., scale-free networks) with small-world attributes shown in **Figure 2m**, a finding that we incidentally previously also postulated for a transcriptomic network of adult hippocampal neurogenesis^21^. The ENR network, however, displayed a heavier tail with a more significant number of densely linked hubs compared to the SD network. As the skewed degree distribution did not become visible in the linear plots, we used logarithmic fitting to present the distributions in a log-log plot (**insets** in **Figure 2m**).

To further evaluate the tolerance against failures and damage, we compared the robustness of SD and ENR scale-free networks by quantifying the changes in the remaining links and node degree when nodes were randomly removed (**Figure 2n**). ENR networks demonstrated higher robustness to random failures of the underlying functional organization than SD networks. In ENR, the communication in the network only slightly changed under increasing errors; even when 100% of the nodes in the SD network were randomly removed, the communication between the remaining nodes in the ENR network was unaffected. This behavior is rooted in the scale-free network topology, in which the majority of nodes have few links. Removal of these nodes did not alter the architecture of the remaining nodes and had no impact on the global network robustness in ENR.

Scale-free networks are known to display failure tolerance, but this comes at the cost of increased attack vulnerability when the highly connected hubs are hit^20^. We thus next quantified the remaining links after removing the highly connected nodes. ENR networks maintained network redundancy for a much higher number of removed nodes before reaching a total loss of communication between links (i.e., the mean lifetime decay of the degree distribution was shorter in the SD networks (25.5 ± 0.124) compared to (96.09 ± 0.33) in ENR) (**Supplementary Figure 3**).

In sum, this section of results illustrated how experience results in large-scale changes in the local and extended hippo-cortical network, expressed in its connectivity and dimensionality^15,16,22,23^. ENR networks exhibited a more pronounced scale-free architecture that renders them a high level of complexity, facilitating functional segregation and global integration, which in turn play a key role in computational power, efficient neural information processing, and promoting network resilience to insult.

### ENR facilitates spatiotemporal firing patterns and increases circuit dynamics

Systematically analyzing patterns of firing synchrony helps to understand the enhanced functional connectivity in our ENR networks. In contrast to conventional small-scale electrophysiological methods such as patch-clamping individual neurons, recordings from large neuronal ensembles provide insights into neuronal synchrony and large-scale network dynamics^24,25^. We thereby mapped the synchronous spatiotemporally localized and coordinated LFP events and multi-unit discharges under SD and ENR conditions. Rhythmic firing initiated in CA3 and propagated to other hippo-cortical subnetworks emerged as slow oscillations (1-100 Hz) superimposed by fast oscillations (<200 Hz). Compared to SD, ENR networks yielded higher synchronous patterns, predominant slow oscillatory activity (i.e., theta band), and increased superimposed ripple-like patterns in CA3, DG, and EC layers, indicated by significantly higher frequency and greater magnitudes in power (**Figures 3a-h**). These features endow the active neuronal ensembles in ENR networks with more rhythm generators^26^ and functionally connected anatomical subnetworks^27^. These arguably support the improved spatiotemporal information coding/encoding, memory consolidation, and synaptic strength^28^ observed under ENR conditions.

**Figure 3.**
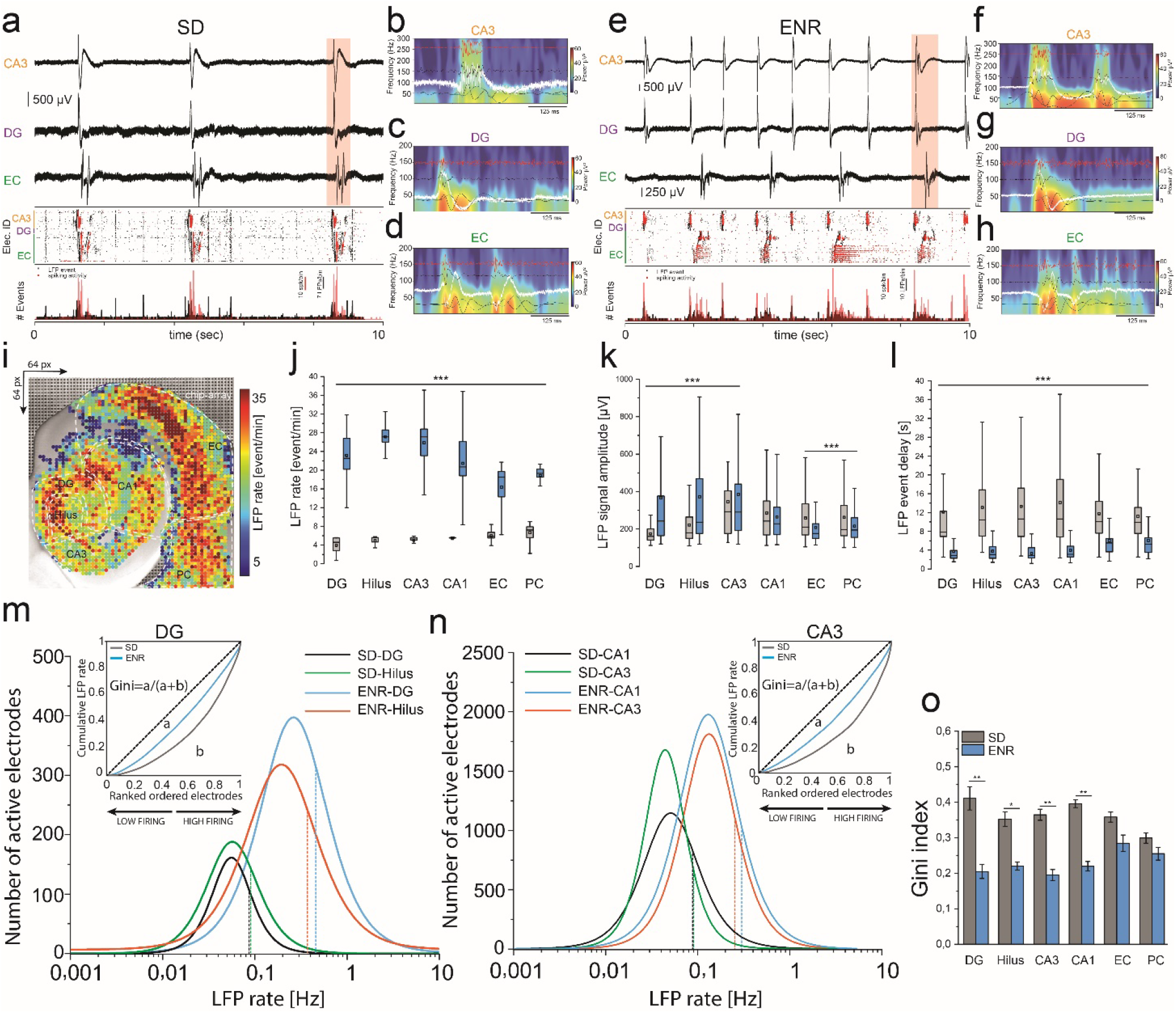
ENR enhances large-scale spatiotemporal firing features and network synchrony. **a)** Representative traces of firing waveforms, aligned with rastergram, and event counts of CA3, DG, and EC layers in SD networks. Rastergram and events count display the interconnected subnetwork with synchronous events (LFPs-black dots and spikes-red). **b-d)** Pseudo-color spectrograms showing the frequency-time dynamics in CA3, DG, and EC firing clusters identified in a, respectively overlaid with bandpass filtered waveforms to identify firing spectrum from theta to ripples in SD. **e)** Same as in a, but from ENR with higher synchronous and frequent events. **f-h)** similar to b-d, but from ENR firing clusters that show a higher magnitude of the theta oscillation and ripple-like activity. **i)** Topographical pseudo-color mapping of ENR large-scale LFP firing pattern overlaid on the hippo-cortical structure. **j-l)** Statistical quantification of the spatiotemporal firing patterns recorded simultaneously from SD vs. ENR networks illustrates significantly higher LFP rates, and amplitude, and faster LFP events (i.e., shorter delay) in ENR compared to SD networks (\*\*\**p < 0.001 ANOVA*) (n=48 slices from 6 mice of SD and ENR each). **m, n)** Firing rate distribution in different hippocampal subregions are skewed in SD and ENR networks and conforming to a lognormal distribution. The distribution in ENR depicts the activation of substantial cell fractions with higher firing rates than SD and contributes significantly to the enhanced neuronal interactions. Dashed lines indicate medians (DG, *p < 10^-100^*, *Hilus*, *p < 10^-137^*, CA3, *p < 10^-14^*, CA1, *p < 10^-21^ Kolmogorov-Smirnov test*). Lorenz statistics (insets) illustrate the neuronal participation in different hippocampal subnetworks (i.e., DG and CA3). **p)** Gini coefficients in different hippo-cortical regions in SD and ENR networks (\*\**p < 0.01, *p < 0.05 ANOVA*).

We next quantified the first-order statistical parameters of the spatiotemporal population firing patterns that demonstrated substantial responses in the ENR subnetworks compared to SD indicated by higher LFP rates, signal amplitude, and shorter delay features between consecutive LFP events (**Figures 3i-l**).

Additionally, we found that in ENR, the neuronal participation in synchronous network events exhibited strongly skewed lognormal distributions with only a small fraction of firing electrodes generating higher firing frequency than SD (**Figures 3m** and **n**, and **Supplementary figure 4**). Compared to SD, the asymmetric firing distributions in the different hippocampal subregions showed heavier tails toward lower frequencies and a significant extension toward higher frequencies in ENR networks (DG, *p <10^-100^*, Hilus, *p <10^-100^*, CA1, *p <10^-21^*, CA3, *p <10^-14^*, EC, *p <10^-93^*, PC *p <10^-27^ Kolmogorov-Smirnov test*).

To further quantify the inequality in the contribution of the firing electrodes across the entire network, we computed the Gini index from the Lorenz curve^29^ (**Figures 3m** and **n insets**): the higher the index, the more unequal is the participation of the firing electrodes. Intriguingly, we observed higher Gini coefficients in SD firing populations (higher inequality with most electrodes firing at a low rate) than in ENR network with many more electrodes contributing to recorded events (**Figure 3o**). In turn, in ENR, the interconnected hippocampal subnetworks of highly firing neuronal ensembles support collective stability and flexibility for continuous network remodeling under rich experience-dependent plasticity without affecting the global functional stability^30^.

In sum, our results identified large-scale functional implications of enhanced oscillatory patterns in hippo-cortical subnetworks of ENR networks. Also, they displayed quantitative and qualitative network electrophysiological differences between SD and ENR networks that opt for higher synchronous dynamics in ENR to support the significant involvement of enhanced ENR subnetwork wiring in rhythm generation and network stability compared to SD^31,32^.

### ENR networks mediate efficient spatiotemporal pattern separation

To link the high-dimensional coding represented by the spatiotemporal firing patterns in the ENR network to concrete network functionality, we next investigated the paradigm of ‘pattern separation’. The hippocampus plays a key role in extracting regularities across experiences from overlapping mnemonic representations that are encoded in the firing of neuronal populations^33^. Previous studies, for example, reported a role of adult hippocampal neurogenesis, which life-long adds new neurons to the dentate gyrus in an activity-dependent manner, in improving the indexing of representations in the DG network under ENR conditions by modulating their firing rates^34^. We here determined the similarity between neural temporal firing patterns in hippocampal subregions on our high-density chip arrays to address the link between input-output transformation and pattern separation mediated by rate mapping^35^.

We defined input and output information as epochs of firing events in EC, DG, and CA3 with simultaneous inter-regional firing and sequential propagation between these regions (i.e., 500, 60, and 100 firing electrodes in EC, DG, CA3, respectively). These interconnected firing sites allowed us to compute the pairwise Pearson’s correlation coefficient (PCC) to determine the similarity between input PCC_(EC-DG)_ and output PCC_(DG-CA3)_, as illustrated in (**Figures 4a** and **b**). In contrast to SD, we found that in ENR, the similarly (correlated) firing events (EC-DG) that are processed in the DG network yielded less similar (decorrelated) output events (DG-CA3) projected to the CA3 (**Figure 4c**). These features were augmented by a “winnertake-all” effect in a computational simulation (see methods), highlighting the criticality of DG network size and the level of firing activity to reduce similar information interference and induce orthogonalization^35,36^. The fact that ENR increases adult hippocampal neurogenesis^37^ suggests a mechanism to fine-tune the size of the functionally relevant neuronal population in the DG^22^, but this aspect was not yet explicitly examined in this study.

**Figure 4.**
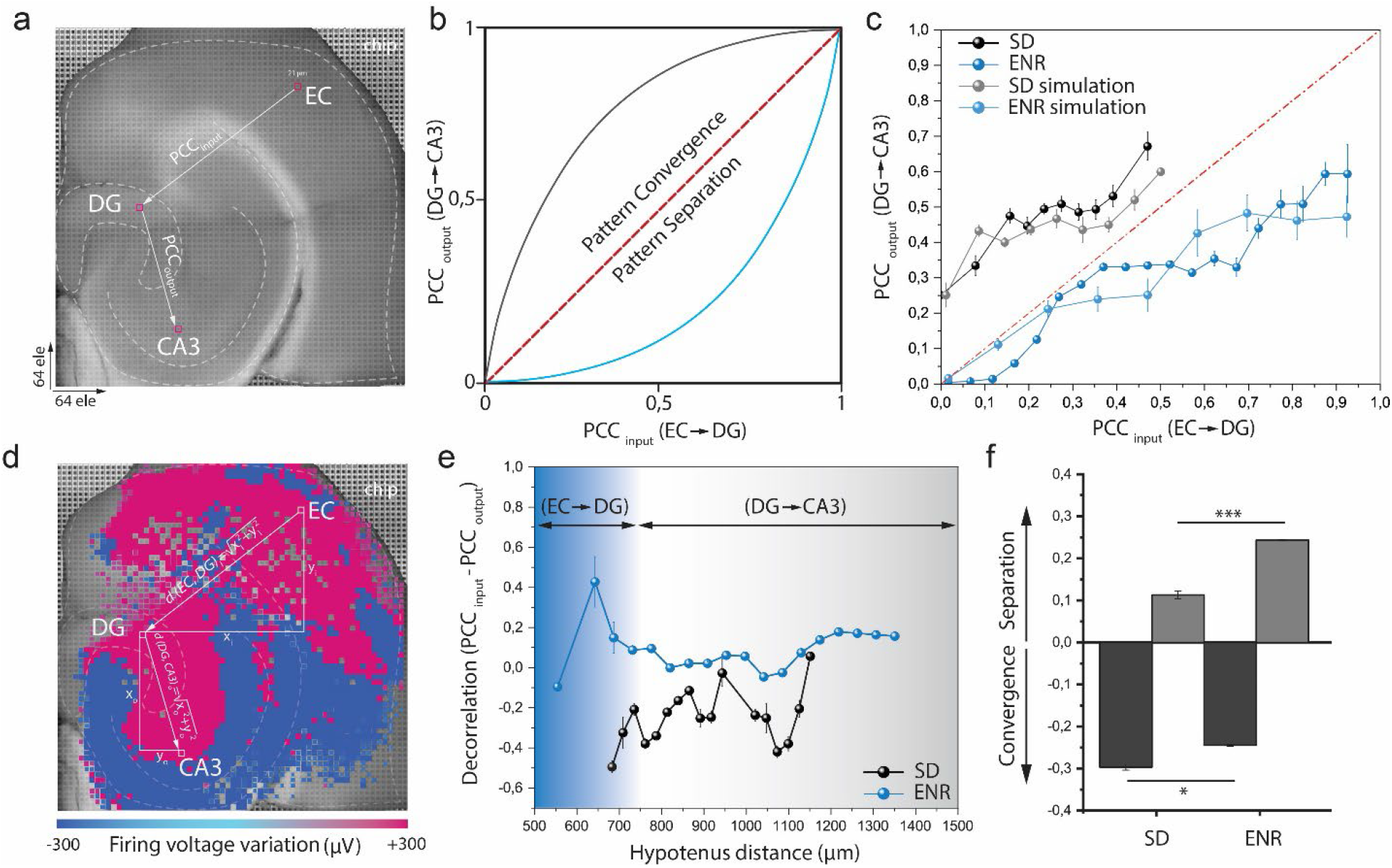
ENR facilitates spatially resolved pattern separation. **a)** Representative image on computing large-scale on-chip input-output firing transformation between (EC-DG-CA3 layers). PCC_input_ and PCC_output_ are the pairwise correlation coefficients for firing electrodes with causal connections between EC→DG and DG→CA3, respectively. **b)** Schematic diagram for scenarios of input-output nonlinear transformation in hippo-cortical subnetworks. When similar input patterns PCC_input_ on x-axis showing lower PCC_output_ the DG network perform a separation (blue line), and convergence at CA3 network when the y-axis is showing higher PCC_output_. **c)** Mean of data points from active firing units corresponding to SD (EC=2387, DG=85, CA3=1583) above the defined dashed red line (convergence), and ENR (*EC*=*3170*, *DG*=*655*, *CA3*=*1804*) below the line (separation) (\*\**p < 0.01 ANOVA*). In ENR (blue), PCC_output_ is lower than PCC_input_, thus indicating pattern separation, whereas SD (black) shows a higher tendency to pattern convergence. The data points are also simulated using a winner-take-all model featuring parameters set automatically from the experimental data points. (\*\**p < 0.01 ANOVA*). **d)** Topographical representation of the emerging input-out pattern transformation with Euclidean distance concept overlaid on the spatial map of the mean-field oscillatory potential. **e)** Mean Euclidean distance of firing electrodes’ spatial representations from EC→DG→CA3. The x-axis indicates the Euclidean distance on-chip computed from the input firing electrode in the EC region, and the y-axis indicates the difference between input-output Pearson’s correlation coefficients. ENR network showing significant decorrelation of the input firing patterns compared to SD network achieved at the DG network (i.e., 500-750 μm on-chip) (*p < 10^-21^ Kolmogorov-Smirnov test*). **f)** Summary of pattern separation and convergence in SD and ENR groups (\*\*\**p < 0.001 ANOVA).*

We next computed the decorrelation between input and output firing patterns as a function of distance. By estimating the Euclidean distance from the input to the output firing sites, we found that the ENR network exhibited a stronger decorrelation and decayed with distance within the dimension of the DG network (500-750 μm), but still remained higher than the SD network (**Figures 4d-f**).

This demonstrated evidence for experience-induced functional modifications in the DG network. This functional modification of the ENR network comprises increased high-dimensional coding and functional connectivity that might enhance pattern separation by providing larger activity space (i.e., decreased noise and variability) to embed different representations with minimum overlap^38^.

### ENR modulates propagation and transmission of firing patterns

A reliable propagation of neural activity is a key prerequisite for carrying information between cascades of active neuronal groups in modular brain circuits^39^. The hippo-cortical circuit processes information in the transverse direction primarily through chemical synaptic transmission mediated by two propagation pathways: serially along the tri-synaptic transmission from EC to CA1^40^ and via recurrent di-synaptic networks between CA3 and DG^41^. Additionally, neural activity propagation at high-frequency oscillations could be mediated by non-chemical transmission using electrotonic coupling via gap junctions^42^, while the propagation of slow oscillatory activity suggested being modulated by non-synaptic ephaptic coupling of the endogenous electrical fields in the neuronal extracellular space^43^. Our multisite recordings across the entire hippo-cortical network allowed quantifying the incidence of spontaneously induced firing oscillations by 4-AP and the spatiotemporal patterns of their long-range propagation. In region CA3 of ENR hippocampi we found a greater incidence of ignition sites of localized firing activity than in SD (**Figures 5a** and **b**). The restricted activity distribution in CA3 propagated broadly towards adjacent and distal areas of CA1 and via the recurrent network pathway to the hilus and DG. Also, an increased probability of activity incidence was prominently found in CA3, CA1, and the EC and significantly higher in all subregions of ENR compared to SD networks (**Figure 5c**).

**Figure 5.**
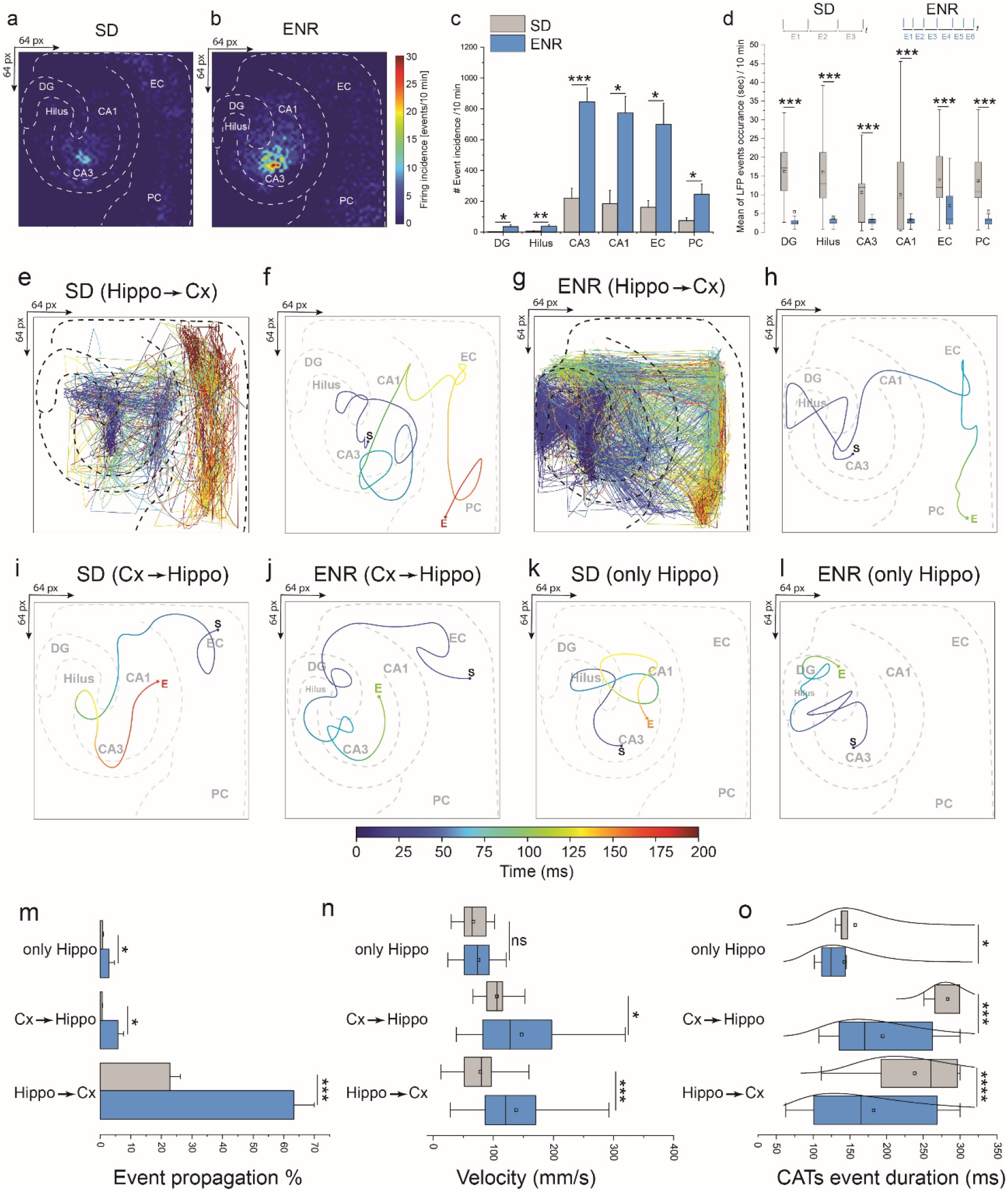
ENR accelerates propagation and modulates firing pattern transmission and dynamics. **a, b)** Incidence analysis of large-scale oscillatory activity patterns. A color scale indicating event probability, computed in 10 min recordings in SD and ENR networks. Both maps show defined initiation points in the CA3 and dynamic event propagation in the hippo-cortical circuitry. **c)** Quantification of event incidence shows significantly higher occurrence in ENR hippo-cortical subregions than in SD. (\**p < 0.05, ***p < 0.001, ANOVA*, n=48 slices from 6 mice of SD and ENR each). **d)** Quantification of event synchrony by event delay indicating a shorter mean duration of activity events in all hippo-cortical subnetworks in ENR compared to SD (\*\*\**p < 0.001*, *ANOVA*, n=48 slices from 6 mice of SD and ENR each). **e, f)** Exemplary 2D CATs superimposed onto the hippo-cortical structural network with a color-coded time scale revealing the propagation patterns in SD network from hippo→cortex. (e) indicating all detected activity events, and (f) showing the average event with starting and end spatiotemporal points. **g, h)** Same as in e & f but for ENR network. **i-l)** Same as in f & h but for propagation patterns from cortex→hippo and intra-hippo in SD and ENR networks. **m)** Quantification of the probability of propagation events based on the classified propagation categories (i.e., hippo→cortex, cortex→hippo, and intra-hippo). (\**p < 0.05*, \*\*\**p < 0.001*, *ANOVA*, n=48 slices from 6 mice of SD and ENR each). **n)** Velocity of conduction based on the firing patterns in the classified propagation categories (i.e., hippo→cortex, cortex→hippo, and intra-hippo). (\**p < 0.05*, \*\*\**p < 0.001*, *ANOVA*, n=48 slices from 6 mice of SD and ENR each). **o)** Quantification of detected CATs event duration for the three classified propagation categories shown in m & n. (\**p < 0.05*, \*\*\**p < 0.001*, \*\*\*\**p < 0.0001*, *ANOVA*, n=48 slices from 6 mice of SD and ENR each).

Next, we identified the topographic propagation delay of the spatiotemporal firing patterns by computing the time-delay between sequential occurring events in each hippo-cortical subregions, which yielded greater firing probability and higher synchrony in the ENR compared to the SD networks (**Figure 5d**). We then analyzed the spatiotemporal propagation patterns of the recorded events with an algorithm based on the center of activity trajectories (CATs)^11^ and classified the CATs of network-wide activity with an unsupervised machine learning algorithm (see method). Thereby, we identified three categories of propagation pathways: 1) hippocampus to entorhinal/perirhinal-cortex, 2) intra-hippocampal circuits mediated by the recurrent network, and 3) entorhinal-cortex to hippocampus corresponding to the classical unidirectional tri-synaptic pathway (**Figures 5e-l**). Quantitatively, our recordings yielded a more prominent propagation probability from the hippocampus to the cortex in ENR (63.2% ± 6.7) than SD (22.7% ± 3.3). Despite an in absolute terms much lower presence of the other propagation categories, in relative terms, ENR still showed a 9-fold increase in cortex to hippocampus propagation and a 3.5-fold increase in intra-hippocampal circuits (**Figure 5m**). We also determined the velocity of conduction in the three propagation categories (**Figure 5n**). Nevertheless, compared to SD networks, ENR showed significantly greater mean velocities than SD (138 ± 3.3 mm/s vs. 78.1 ± 2.3 mm/s, hippocampus → cortex, and 147 ± 10 mm/s vs. 105 ± 14.3 mm/s, cortex → hippocampus). These results were further supported by computing the duration of the CAT events in each propagation pathway. We found significantly shorter event duration in all CAT event categories in ENR vs. SD networks (**Figure 5o**).

In sum, these findings demonstrated the impact of rich experience on modulating large-scale neural activity propagation characteristics with a faster information processing in the ENR than the SD networks. This enhancement is mediated by the feedback loop from CA3 to DG, whose sharpened network synchrony is relevant for storing and retrieving information sequences^44^ and facilitating pattern separation^45^. The propagation features in ENR networks might be associated with increasing the hippocampus’s tightly layered organization, thus facilitating stronger endogenous local fields to induce a significant effect on neural synchronization^46^. Nevertheless, the greater conduction velocities of the bidirectional hippocampus ↔ cortex propagation observed in ENR than SD networks might implicate a critical combinatorial mechanism for spatiotemporal propagation in hippo-cortical subregions influencing neuronal functions and dynamics. This mechanism could be facilitated by synaptic mode (via a recurrent loop) and non-synaptic mediated by electrotonic^42^ and ephaptic couplings^43^. However, investigations by large-scale experimental paradigms and computational modeling are further needed to rule out the exact mechanism of activity propagation in ENR.

### Experience boosts neural information encoded in the spike-phase dynamics

The brain encodes information using several coding schemes employed individually or concurrently. Both experimental and theoretical evidence supports the idea that in the hippocampus, information is encoded with temporal coding by theta rhythm phase precession^47,48^. This notion builds on encoding neural information of firing activity (i.e., LFPs and spikes) on different temporal scales to investigate circuit mechanisms underlying the functional connectivity^49^ and neural representations in spatial and non-spatial information^50,51^.

To study whether ENR would also enhance the encoding of higher information compared to SD conveyed by the LFP phase at the time of spiking activity, we used our large-scale extracellular recordings of LFPs and spikes. Thereby, we quantified the relationship between the LFP and multi-unit spiking activity using the “phase-of-firing” coding paradigm and information theory of Panzeri and colleagues^52^.

A first analysis of the LFP spectrum in both ENR and SD indeed revealed the highest power in the oscillatory theta band (4-12 Hz; **Supplementary figure 5a**), so that we considered theta oscillation a reliable frequency range to encode information at all firing electrodes in the network. We then quantified the LFP phases by assigning them to four color-labeled quadrants of equal duration (**Figures 6a** and **b**). We employed the phase-of-firing coding to identify those LFP phase quadrants during which spikes were discharged over long periods (**Figures 6e** and **f**). This allowed us to identify supposedly specific information contents encoded in the spatiotemporal patterned activity as opposed to the less specific spike count code shown in **Figures 6c** and **d**. Indeed, our results suggested richer information content in the spike rate associated with the LFP phase than observed in the spike counts alone (**Figures 6g** and **h**). Information encoded in the LFP phases of all four quadrants was significantly greater in the ENR network than SD (**Figure 6i**) and strongly depended on the LFP frequency range (theta). For all firing electrodes, the strongest preference of phase-associated spikes was in the 0 →π/2 range (the red positive rising phase of the oscillation) followed by the π→ 2π range (the green and blue trough of the oscillation).

**Figure 6.**
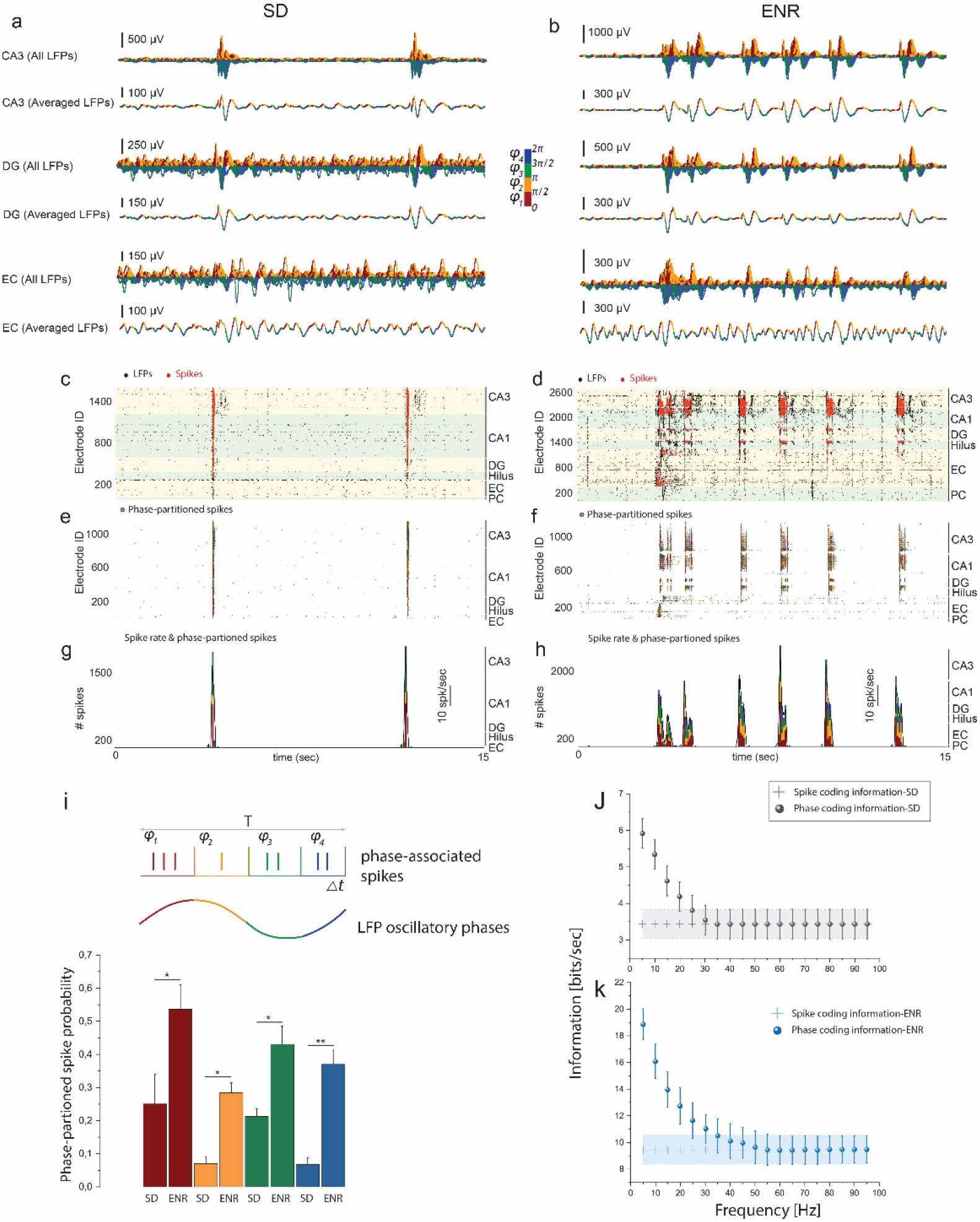
ENR enhances information encoded in the phase-of-LFP oscillation. **A, b)** For SD and ENR networks, waveform traces of LFP activity from large-scale recordings of hippo-cortical layers (CA3, DG, and EC) showing all extracted and averaged waveforms in the theta oscillatory range (4-12 Hz) that are color-coded with four quadrants indicating the associated phase of the LFP firing. **c, d)** Raster plots of SD and ENR networks showing LFP and multiunit spiking activity in all interconnected hippo-cortical subnetworks. Black dots for LFPs and red dots indicate spikes. **e, f)** The same time course in c and d but only for spikes color-coded by the phase of LFPs at which the spikes were emitted. **g, h)** Multiunit spike rate in SD and ENR networks (black lines) and the amount of spikes is computed according to the LFP phase (color-coded), which indicates the same spike rate (SD vs. ENR) encode additional information associated with the LFP phase. **i)** For a given time course of spike trains, the multiunit spiking activity can be partitioned based on the slow oscillatory LFP signal into four color-coded phase quadrants. Thus, the spiking probability in the LFP phases is computed for SD and ENR networks. Compared to SD, the ENR network shows significantly higher information encoded in all quadrants of the firing phases. (\**p < 0.05*, \*\**p < 0.01*, *ANOVA*). **j)** Rate coding vs. phase coding to compare the conveyed information in SD and ENR hippo-cortical firing information. **J)** The phase of firing information in SD. **k)** The phase of firing information in ENR. Plus signs show the spike rate information in SD and ENR networks in j-k.

We further substantiated these findings by quantifying that spike counts alone conveyed 9.45 ± 1.18 bits/s of information in ENR networks compared to 3.43 ± 0.39 bits/s in SD (**Figures 6j** and **k**). Additionally, however, the 4-12 Hz LFP phase carried 5.91 ± 0.40 bits/s in SD, but 18.85 ± 1.16 bits/sec in the ENR network, indicating an over proportional gain in phase-dependent encoding in ENR compared to SD (**Figures 6j** and **k**). The phase of oscillatory activity conveyed 99% in ENR vs. 72% in SD of additional information not obtained from spike counts. Although spike counts encoded more information in ENR than in SD, the theta LFP phases carried prominently much higher information that was not obtained from spike count code alone. With increasing frequencies, the extra information encoded in the LFP phases declined to reach the spike count information at frequencies greater than 25 Hz in SD and 35 Hz in ENR. Thus, information carried by LFP phases of ENR and SD networks at higher frequency bands (13 - 30 Hz and 30 – 100 Hz) was lower compared to 4 −12 Hz but still carried more information than spike counts (**Supplementary figures 5b** and **c**).

In summary, these findings demonstrated that rich experience enhanced the density of encoded information. This was achieved by adding higher-dimensional complementary layers of information consisting of labeling spikes by their relation to the phases of theta oscillation. Such phase coding has been established as a mechanism for spatial representations^47^, where theta oscillations provide a pacemaker for the spike trains that organize spatial information^48^.

Thus, the impact of the enriched experience in improving the characteristics of rhythmic dynamics suggests a mechanistic role for higher information processing during the offline state of animal behavior. Such a mechanism appears relevant not just for spatially modulated ensembles but even for intrinsic neuronal assemblies in the hippo-cortical slices isolated from the rest of the brain in our experiment^53^. Hence, previous ENR experience endowed the network with the capacity to exploit multiplexed encoding to enhance the information dimensionality and foster the stability and robustness of the neural code^54^.

## CONCLUSIONS AND OUTLOOK

In the present study, we used a high-density neurochip platform to obtain next-generation large-scale electrophysiological patterns across hippo-cortical networks from mice reared in standard and enriched environmental housing conditions. We developed a deterministic method that combined neural information from network-wide firing time courses and features of graph theory to unveil the impact of experience on local and global brain functional connectivity. Our large-scale multisite measurements have overcome challenges associated with the unmet need for monitoring the hippo-cortical subregions simultaneously with a high spatiotemporal resolution that allowed quantifying input-output neural transformation, network synchrony, and dynamics. This illustrated a unique link between oscillatory firing patterns and the topological network arrangement under ENR exposure. Thus, our work suggests a new tool to address network-wide plasticity underlying ENR spatiotemporal coding, not accessible with current bioimaging, electrophysiological methods, and computational models. To our knowledge, this is the first demonstration of mesoscale *ex vivo* connectome and spatiotemporal firing patterns modulation in the hippocampus by ENR. Hence, our findings could provide a pathway to discover fundamental mechanisms of experience-dependent enhancement of hippocampal network underlying high brain functions of memory, navigation, and congnition^1,3^.

Indicated by the featured network electrophysiology and rigorous algorithms from network science in ENR compared to SD networks, we showed that ENR enhanced local functional connections to optimize network-wide rewiring, adaptability of functional remodeling, and firing pattern evolvability by preserving the functional integrity of the subregional networks. Further, the enhanced connectome in ENR subnetworks displayed densely linked hub nodes characterized by a scale-free small-world topology that revealed high-dimensional coding, facilitated efficient pattern separation, and enhanced the density of encoded information for faster processing of the propagating neural activity.

Altogether, our study could explain the prevailed computational benefits of a large-scale connectome for spatial information remapping upon neurogenesis-induced experience-dependent enhancement, providing a fundamental mechanism for improved spatial learning and memory-related processes^55,56^. Interestingly, fostered rhythmic dynamics could also explain the impact of enrichment on brain resilience stemming from the cognitive reserve theory and how the hippocampal circuitry takes advantage of oscillatory dynamics and synchrony in information transfer that could be altered in neurological disorders^57^. Prospectively, modulating the network-wide activity using the advent of electrical and optogenetic approaches may also provide a potential tool to interrogate specific subnetworks along with large-scale experience-dependent enhancement in the hippocampal circuit.

## MATERIALS AND METHODS

### Animals and hippocampal acute slice preparation

All experiments were performed on 12-week-old C57BL/6j mice (Charles River Laboratories, Germany) in accordance with the applicable European and national regulations (Tierschutzgesetz) and were approved by the local authority (Landesdirektion Sachsen; 25–5131/476/14). Two groups of mice were used, either housed in a standard (SD) environment or an enriched environment (ENR) for 6 weeks prior to experiments. Mice were anesthetized with isoflurane before decapitation. Brain slices were prepared according to our previous study^11^. Briefly, the brain was carefully removed from the skull and placed in a chilled cutting sucrose solution before slicing. The brain was fixed on the cutting plate, and horizontal slices (300 μm thick) were prepared using Leica Vibratome VT1200S (Leica Microsystems, Germany). Slices were cut at 0–2 °C in aCSF solution saturated with 95% O_2_ and 5% CO_2_ (pH = 7.2–7.4) of a high sucrose solution containing in mM: 250 Sucrose, 10 Glucose, 1.25 NaH_2_PO_4_, 24 NaHCO_3_, 2.5 KCl, 0.5 Ascorbic acid, 4 MgCl_2_, 1.2 MgSO_4_, 0.5 CaCl_2_. Next, hippocampal slices were incubated for 45 min at 34 °C and then allowed to recover for at least 1 hour at room temperature before recordings with a high-density neurochip. Perfusate used during recordings contained in mM: 127 NaCl, 2.5 KCl, 1.25 NaH_2_PO_4_, 24 NaHCO_3_, 25 Glucose, 1.25 MgSO_4_, 2.5 CaCl_2_, and was aerated with 95% O_2_ and 5% CO_2_.

### Large-scale extracellular recordings from hippocampal networks

Extracellular recordings were performed using high-density (HD) CMOS-biosensors and an acquisition system (3Brain AG, Switzerland) customized to our recording setup. HD neurochips are composed of 4096 recording electrodes with a 42 μm pitch size to compose an active sensing area of ~7 mm^2^, ideal for recording entire hippocampal-parahippocampal regions. The on-chip amplification circuit allows for 0.1–5 kHz band-pass filtering conferred by a global gain of 60 dB sufficient to record slow and fast oscillations^11^. Slices were moved and coupled onto the neurochips using a custom-made platinum harp placed above the tissue. A steady perfusion system was built to deliver oxygenated recording perfusate to slice-neurochip interface with a flow rate of 4.5 mL/min and was temperature controlled at 37 °C throughout the experiment and recordings. We collected extracellular recordings at 14 kHz/electrode sampling frequency from spontaneous and pharmacologically evoked activity using 100 μM 4-aminopyridine (4AP) (Sigma-Aldrich, Germany). A modular Stereomicroscope (Leica Microsystems, Germany) was designed and incorporated into the setup to capture the acute slices light-imaging simultaneously with the whole-circuit extracellular recordings. These images were further used to obtain spatial organization of tissue relative to firing electrode layout during analysis. Recording sessions and LFPs and multi-unit activity (MUA) events detection with hard threshold and precise timing spike detection algorithms (PTSD)^58^ algorithms were performed with commercially available software (3Brain AG, Switzerland). Detected events were further processed and filtered with a low-pass filter (1-100 Hz) for LFPs, and a band-pass filter (300-3500 Hz) for MUAs, and a band-pass filter (140 - 220 Hz) for ripples. Additionally, LFP waveforms were bandpass filtered using 4^th^ order Butterworth filter at delta (1-4 Hz), theta (5-12), beta (13-35), and gamma (35-100) bands.

### Data Analysis

All basic and advanced algorithms used in this work were developed and implemented with custom-written Python scripts. Any package add-ons are cited accordingly.

### Structural clusters

To characterize hippocampal network behavior locally and globally, firing electrodes were structurally related to a specific hippocampal region. Thus, light microscope hippocampal images were overlayed on the neurochip microelectrodes layout. Electrodes were then grouped into clusters based on structural markers on the hippocampal slice. These clusters included six major regions of the hippocampal-parahippocampal circuit −DG, hilus, CA1, CA3, EC, and PC.

### Functional connectivity and causality

To infer large-scale statistical dependent connectivity on a multilayered hippocampal network, cross-covariance was calculated between pairs of electrodes in the 64 x 64 array using Pearson’s correlation coefficient (PCC). This was followed by performing Multivariate Granger causality to quantify the influence of one time series on another and Directed Transfer Function (DTF) to measure directional information flow as we previously described^11^. All the electrodes were sorted by clusters, and the statistic calculations for PCC and interconnection links count were based on paired clusters.

### Network connectivity metrics

Graph Theory was used to characterize overall network topology and interconnectedness from detected LFP events. We computed the graph metrics in custom-written Python code and adapted functions from NetworkX-python package^59^, available on GitHub (https://github.com/networkx). Briefly, the functional network connectivity metrics are described by considering the node *n* as the central component of the graph that may or may not be connected to one another. In our case, a node corresponds to a specific electrode in the sensing array. Also, the edges *e* are the functional links or connections between each node *n*.

#### Degree

To characterize the different representations of network connectivity, we characterized the degree *k* of a node *n* to describe the number of edges connected to a node. We also computed the in-degree or out-degree based on the direction of the vectors (i.e., the flow of information).

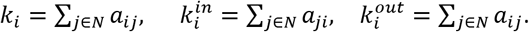

Where *k_i_* denotes the degree of a node *i. a_ij_* denotes the connection between nodes *i* and *j. N* is the set of all computed nodes in the network, 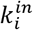 is in-degree and 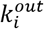 out-degree of node *i*, respectively.

#### Degree centrality

To determine nodes with high topological centrality and influence on network function, the centrality (*DC_i_*) was computed as the normalized fraction of nodes that node *i* connected to^13^.

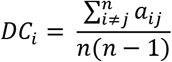

#### Clustering coefficient

To measure how nodes in a given network tend to cluster together to assess the functional segregation, the clustering coefficient (*CC_i_*) was computed^13^.

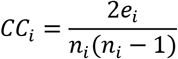

Where *n_i_* denotes the number neighbors to node *i*, and *e_i_* denotes the number of links connecting the *n_i_* to the node *i*.

#### Transitivity

To reveal the existence of tightly connected communities, the transitivity was characterized as the fraction of all possible connected triangles (*N_triads_*) to describe two edges with a shared vertex in a network^13^.

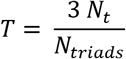

Where *N_t_* is the actual number of triangles in a network and *N_triads_* is the number of triads or possible triangles that consist of two edges with a shared vertex.

#### Average shortest path length

To measure the functional integration of the information flow between layers in the network, the average shortest path length (*L_i_*) was computed^13^.

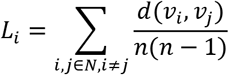

Where *d*(*v_i_*, *v_j_*) denotes the shortest path from *i* to *j*, and is normalized over all possible paired number of nodes *n* in the network.

#### Hub nodes and rich club nodes

To determine centralized, important nodes in a network and reveal network topology, hub nodes and rich club nodes were analyzed. Hub nodes were detected based on three nodal metrics - node strength, clustering coefficient, and network efficiency. The metric value for each node was calculated and compared to determine whether the node value was in the top 20% of all nodes^60^. To restrict the definition of the hub node, we set limits with a hub score. Our hub score was valued between 0 and 3, where nodes either satisfied the top 20% in none, 1, 2, or all 3 nodal metrics. Within the hub node group is a subgroup of nodes with dense connections that conferred the rich-club nodes and are described as hub nodes with a higher degree than the average and provided by the rich club coefficient *ϕ*(*k*)^61^.

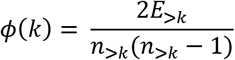

Where *k* denotes the degree, *n*_>*k*_ represents the number of nodes whose degree is larger than a given value *k*, and *E*_>*k*_ denotes the number of connections in a subnetwork comprising *n*_>*k*_.

### Network topology characterization

To determine the potential impact of hub nodes on the network function and the organizational processes shaping network topology, we characterized the degree distributions *P*(*k*) of detected nodes in SD and ENR datasets, which resulted in decayed distribution with a power-low tail^19^.

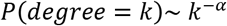

To estimate the power-low degree distribution *P*(*k*) to describe the scale-free topology with a small-world attributes, we used the lognormal model fit.

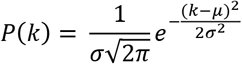

Where *μ* and *σ* are the mean and standard deviation of the distribution, respectively.

To visualize the best-fit network characterization, a complementary cumulative distribution function (cCDF) was used instead of the probability density of the node degree and plotted on logarithmic axes for a more robust visualization for the high-k regime. Goodness-of-fit tests were performed between actual data and fitted models and were estimated by the coefficient of determination *R*^2^.

### Robustness

To estimate the robustness and flexibility of networks for error tolerance and attack vulnerability, we employed two strategies; i) for error tolerance, we measured the node degree and remaining links after a fraction of nodes were randomly removed; ii) for attack vulnerability, we measured the remaining links in network after stepwise removal of high degree nodes.

### Graph map visualization

To visualize the large-scale network connectivity maps, we constructed the data architecture containing nodes and edges. This setup allowed marking the node IDs, coordinates, labels, edge sources, and edge targets. These maps were then converted into (.gexf) file format and were directly read and visualized in the Gephi program 9.2 version (https://gephi.org). To compare the ENR vs. SD, the top 2% of the total functional links were plotted with similar edge weights and degree range queries.

### Dimensionality

To characterize the link between network-wide collective activity and connectivity, we computed network dimensionality^16^. The functional connections or links between paired electrodes across all LFP events in a recording time window served as an input. The dimensionality is defined as the weighted measure of the number of axes of all firing electrodes in the activity population space.

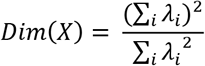

Where *λ_i_* is the *i^th^* eigenvalue of covariance matrix *X*. The resulting value was normalized over the maximum dimensionality value to give a data range from 0 to 1.

### Time-frequency and power spectrum density (PSD)

To determine the power distribution of the LFPs and the dominant frequencies within an oscillatory event, we constructed a frequency-time dynamic in pseudo-color spectrograms for a selected time window using filtered LFPs (1-100 Hz). To estimate the PSD, we used Welch’s method as previously described^11^ by calculating the Fast Fourier Transform (FFT) of the detected LFPs.

### Mean activity basic analysis

We selected three parameters to describe the mean activity features of large-scale spatiotemporal LFP oscillations, including LFP rate, amplitude, and event delay. LFP rate was defined as the number of detected LFP events per minute, and the delay was defined as the time between detected events. The signal amplitude analysis was obtained through full-wave rectification and low-pass filtering with a cut-off frequency of 100 Hz.

### Lognormal distribution

The LFP firing patterns of the hippocampal neuronal ensembles showed a wide degree of participation in the circuit activity and followed a skewed lognormal distribution. Thus, we computed the probability density function for the lognormal distribution as previously described^11^.

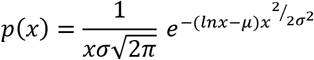

Where *μ* is the mean, and *σ* is the standard deviation.

### Gini coefficient

To quantify the inequality of participation of individual ensembles in the interconnected layers of the hippocampal circuit, we employed the Gini index, which was computed as the ratio of the areas on the Lorenz curve diagram^11^.

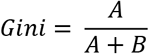

Where *A* is the area above the Lorenz curve, and *B* is the area below for the cumulative LFP firing rates.

### Large-scale pattern separation

To determine the similarity between neural temporal firing patterns to address the link between input-output transformation and pattern separation, we assessed the similarity between input patterns of firing activity from EC to DG and output patterns from DG to CA3. All active electrodes with functional causal connections in the EC-DG-CA3 pathway were selected. Therefore, we defined the positions for input sources in the EC, pattern separators in the DG, and target sites in the CA3, which were associated with the same firing electrodes in the DG. Similarity between pairs of electrodes was based on the waveforms in the related LFP event timescale and was computed using Pearson’s cross-correlation. For a given pair of variables (*X_input_*, *Y_output_*), correlation coefficient values were calculated for input and output regions *PCC_input_* and *PCC_output_* that are contained in the set {***PCC**_input, output_*} ⊇ ({*PCC_input_*},{*PCC_output_*}) which could be defined as,

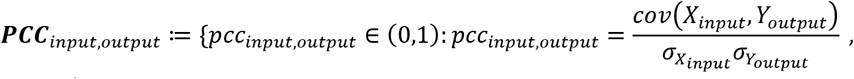

Where (*X_input_*, *Y_output_*) are from input and output electrode sites, respectively, and *σ* is the standard deviation.

The relationship between pairs of electrodes was classified as pattern separation when *PCC_output_* was lower than *PCC_input_*, and pattern completion when *PCC_output_* was higher than *PCC_input_*. The decorrelation was defined as the difference between the covariance of input and output pairs of firing electrodes, *PCC_input_*, and *PCC_output_*, respectively. Further, to quantify the relationship between the decorrelated input and output and the distance from the DG layer, we computed the Euclidean distance *D* based on the positions of input and output electrodes and formulated as,

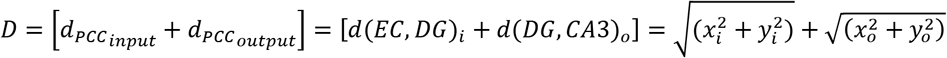

Where, the pathway distances {*x_i_*, *y_i_*} are computed only between active electrodes in the input region, and similarly {*x_o_*, *y_o_*} are the pathway distances between the active electrodes in the output region.

Next, to illustrate how the size of the DG layer influences the process of pattern separation and completion, we implemented a simplified computational model for the trisynaptic pathway EC-DG-CA3 without considering the recurrent dynamics in the CA3, and we used a k-winner-take-all algorithm for selecting the active electrodes^62^ (i.e., sparse activation). The network was built with three-layer architecture, where the EC and CA3 represented the input and output layers, respectively, and the DG as a hidden layer. We set the input layer parameters, including the size of the DG layer indicated by the number of active electrodes, the *PCC_input_*, and the *PCC_output_* ranges. Varied sizes of the DG layer were set while other input values were generated randomly with multi-dimensional Gaussian distributions under the constraints of the value ranges, which were obtained from the empirical datasets. The networks were trained to recode their inputs to the output layers at varied sizes of the DG to test the impact of adding new active units in the DG layers upon neurogenesis in the enriched environment. Finally, to illustrate pattern separation and completion comparison in SD and ENR networks, we calculated the differences between input and output cross-correlation coefficient values for DG size of 50, and 300 active electrodes in SD and ENR networks, respectively.

### Spatiotemporal LFP tranjectories

To quantify the propagation magnitude of the spatiotemporal LFP events in all interconnected hippocampal layers, we employed the center-of-activity trajectories (CATs) analysis^11^. Voltage values embedded in the LFP frame activities within a 5 ms moving time bin were used to collect the CAT magnitudes in each firing event. The value of the CAT at time *t* is a two-dimensional vector.

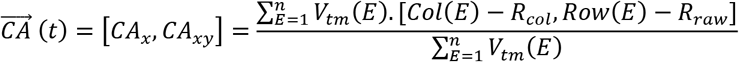

Where *V_tm_*(*E*) denotes the LFP firing rate corresponding to the active electrodes *E* within a time window (*tm*). *Col*(*E*) and *Row*(*E*) are the column and row numbers of the associated *E. R_col_* and *R_row_* are the coordinates of the physical center of 64 x 64 electrodes. *n* is the total number of active electrodes. Then, the CA trajectory from *t_0_* to *t_1_* with a time step Δ*t* can be computed.

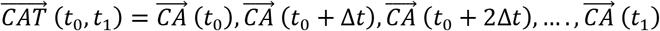

Based on the spatiotemporal trajectories for all the firing events, we categorized the propagation pathways of firing events in the interconnected hippocampal-parahippocampal network into three major groups; i) from the hippocampus to entorhinal-perirhinal regions (Hippo →EC-PC), from entorhinal regions to hippocampus (EC-PC →Hippo), and hippocampus only (Only Hippo).

### Velocity of conduction

To track the putative displacement of LFP events over the entire hippocampal circuit, the velocity of conduction was calculated as the average of instantaneous vector quantity displacements calculated for the time of a propagating LFP event^11^.

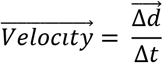

Where Δ*d* is the change in displacement and Δ*t* is the change in time.

### LFP-spike information analysis

To quantify the information carried by phases of individual LFP fluctuations at the time of multi-unit spiking activity, we employed a four-step procedure; i) we extracted individual frequency information from a multilayered hippocampal-parahippocampal network at (4-12 Hz) band as resulted from our power frequency analysis; ii) we computed the instantaneous phase values (i.e., with *π*/2 precision, where the phase values 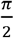 and 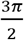 correspond to the peak and trough of the oscillation, respectively) of the narrow-band waveform fluctuations in multiple events using the Hilbert transform. This allowed visualizing the LFP phases in color-coded equispaced quadrants; iii) we extracted multi-unit spikes (MUA) from the interconnected hippocampal layers using the precise timing spike detection algorithm^58^ with a threshold set to 8 standard deviations of the band-pass filterd signal (Butterworth filter, 4^th^ order) at (300 – 3500 Hz). We then labeled the spikes with the color of the LFP phase quadrants at the time of their occurrence and presented them in rastergrams and rate code plots; iv) next, we quantified the information carried by the phase of the LFP using the Shannon theory^63^. Since the observation value series is continuous^64^, the information *I*(*T*) is characterized in units of bits, carried by LFPs and MUA in time window *T*;

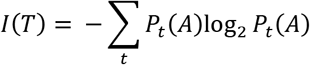

Where *P_t_*(*A*) is the probability of observing LFP or MUA event series across the time window of recordings.

### Statistical analysis

All statistical analyses were performed with Originlab 2020. All data in this work were expressed as the mean ± standard error of the mean (SEM). All box charts are determined by the 25^th^-75^th^ percentiles and the whiskers by the 5^th^-95^th^ percentiles and lengths within the Interquartile range (1.5 IQR). Also, the lines depict the median and the squares for the mean values. Differences between groups were examined for statistical significance, where appropriate, using Kolmogorov-Smirnov test, one-way analysis of variance (ANOVA), or two-way ANOVA followed by Tukey’s posthoc testing. P < 0.05 was considered significant.

## ACKNOWLEDGMENT

We would like to thank the DZNE within the Helmholtz Association for funding this study. We would also like to acknowledge the platform for behavioral animal testing at the DZNE-Dresden (Alexander Garthe, Anne Karasinsky, and Sandra Günther) for their support.

## AUTHOR CONTRIBUTIONS

BAE: Performed part of experiments and analyzed the data.

XH: Wrote the code for analysis and analyzed the data.

SK: Performed part of experiments and analyzed the data.

GK: Conceptualized project, designed and performed animal experiments, supported interpretation of results, provided co-funding, co-wrote the manuscript.

HA: Conceptualized project, planned and managed the project, designed and performed experiments, analyzed the data, developed analytical tools, and co-wrote the manuscript.

All authors revised, reviewed, and approved the final version of the manuscript.

