## Supplementary Figures for "Rich Experience Boosts Functional Connectome and High-Dimensional Coding in Hippocampal Network"

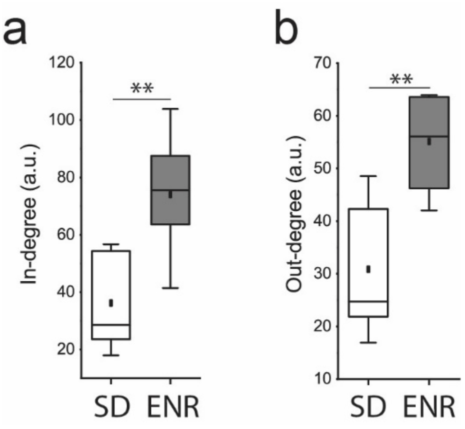


**Supplementary Figure 1 |** ENR network exhibits higher complexity shown by **a)** in-degree and **b)**out-degree metrics *(**p < 0.01 ANOVA*).

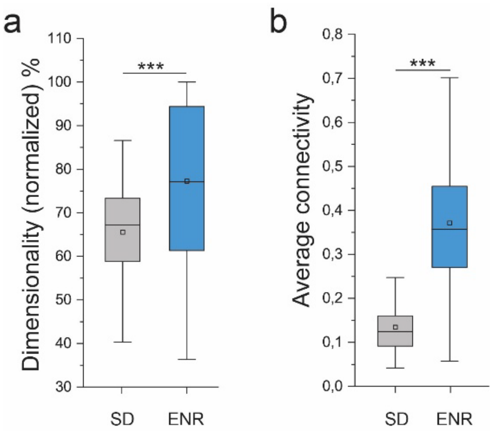


**Supplementary Figure 2|** Quantification of network dimensionality depicts higher dimensional coding in ENR than SD, indicated by **a)** normalized percentage of dimensionality and **b)** network dimensionality as a function of average connectivity *(***p < 0.001 ANOVA*).

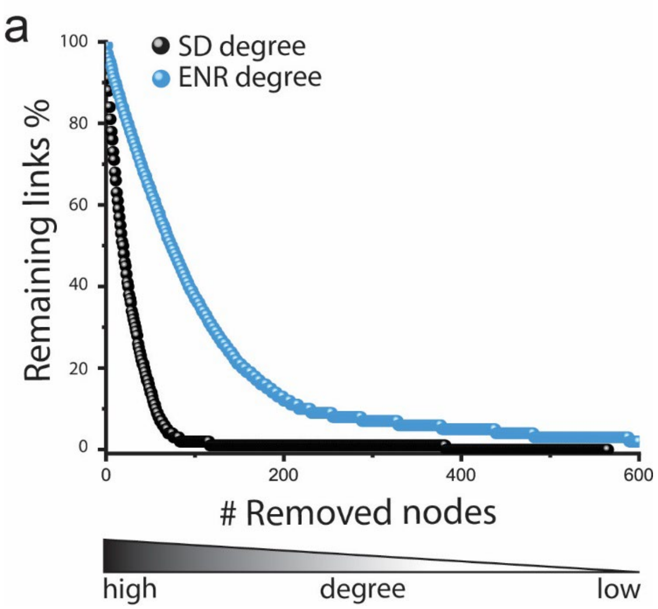


**Supplementary Figure 3|** ENR networks display higher redundancy and failure tolerance than SD quantified by the percentage of remaining links after removing nodes from the highest to the lowest degree. The mean lifetime decay of the degree distribution is shorter in the SD networks (25.5 ± 0.124) compared to (96.09 ± 0.33) in ENR *($p < 10^{-10}$ Kolmogorov-Smirnov test*).

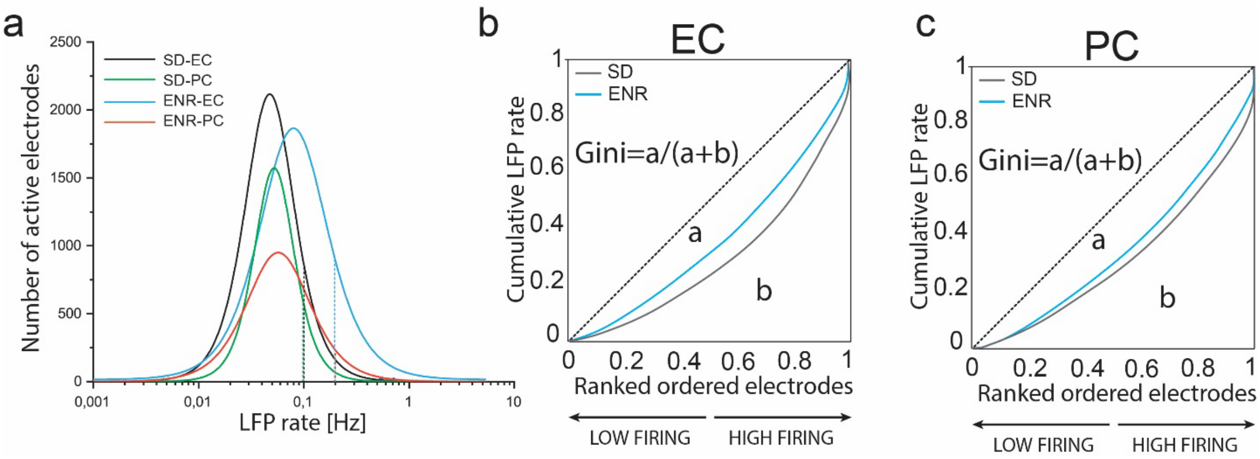


**Supplementary Figure 4|** Lognormal distribution and Gini analysis. **a)** Firing rate distribution in EC and PC subnetworks are skewed in SD and ENR and conforms to a lognormal distribution. Dashed lines indicate medians (EC, $p < 10^{-23}$, and PC, $p < 10^{-27}$ *Kolmogorov-Smirnov test*). **b-c)** Lorenz curves illustrate the neuronal participation in EC and PC layers.

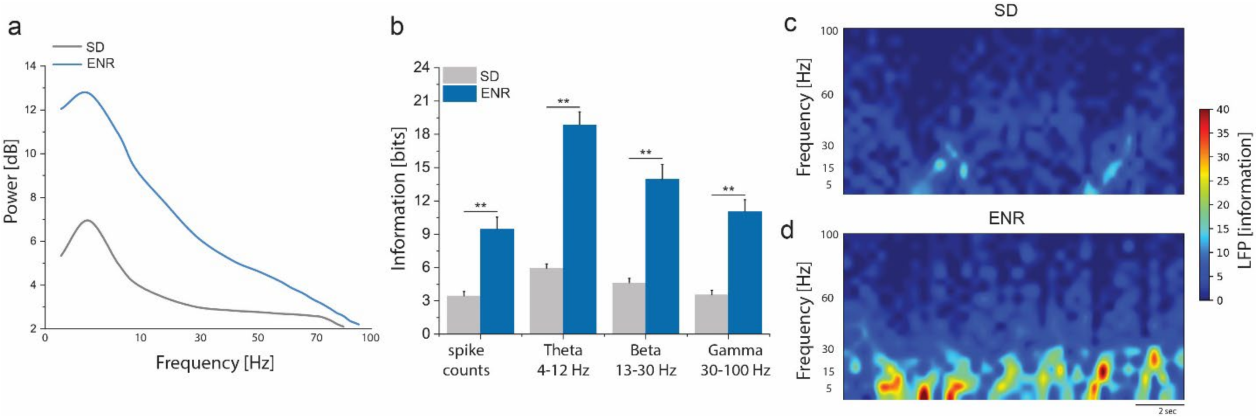


**Supplementary Figure 5| a)** Spectral features of LFPs in the hippo-cortical recordings showing an increase of power magnitude at low-frequency theta band (4-12 Hz), albeit higher in ENR than SD (*p < 0.05 Kolmogorov-Smirnov test*. **b)** ENR showing a higher density of Information rates than SD conveyed by spike counts and the LFP phase at which spikes were discharged as a function of three LFP-frequency bands *(**p < 0.01 ANOVA*). **c)** Pseudo-color spectrograms showing higher LFP information in ENR than SD conveyed by low oscillatory frequency theta rhythm (4-12 Hz).
